# Cadmium interference with iron sensing reveals transcriptional programs sensitive and insensitive to reactive oxygen species

**DOI:** 10.1101/2020.07.05.188649

**Authors:** Samuel A. McInturf, Mather A. Khan, Arun Gokul, Norma A. Castro-Guerrero, Ricarda Hoehner, Jiamei Li, Henri Margault, Hans Henning Kunz, Fiona L. Goggin, Marshall Keyster, Rachel Nechushtai, Ron Mittler, David G. Mendoza-Cózatl

## Abstract

Iron (Fe) is an essential micronutrient whose uptake is tightly regulated to prevent either deficiency or oxidative stress. Cadmium (Cd) is a non-essential heavy metal that induces both Fe-deficiency and oxidative stress; however, the mechanisms underlying these Cd-induced responses are still elusive. Here we explored Cd-induced Fe-associated responses in wildtype *Arabidopsis* and *opt3-2*, a mutant that over-accumulates Fe. Gene expression profiling revealed a large overlap between transcripts induced by Fe-deficiency and Cd exposure in wildtype plants and the *opt3* mutant. Interestingly, vascular-localized Fe-responsive genes were found to be highly induced by Cd even in the presence of high Fe and H_2_O_2_ levels, suggesting that Cd impairs Fe sensing. It was recently shown that Fe-S cluster-containing proteins AtNEET, play a role in Fe sensing. Our data shows that Cd negatively impacts both the stability and Fe-S transfer activity of AtNEET. Altogether, our data indicate that Fe-deficiency responses are governed by multiple inputs and that a hierarchical regulation of Fe-deficiency responses prevents the induction of specific gene networks when Fe and H_2_O_2_ levels are high. Other Cd/Fe-responsive genes however, are insensitive to this negative feedback regulation suggesting that their induction is the result of an impaired Fe sensing as opposed to the traditional view of Cd/Fe uptake competition at the root level.

**Highlight:** Cadmium induces an iron-deficiency response often explained by root uptake competition; here we show that Cd also impairs Fe sensing in leaves, even when Fe is in sufficient quantities.

## Introduction

Iron (Fe) is an essential nutrient for all known biological systems, facilitating the transfer of electrons and acting as a cofactor in metalloproteins. While vital for life, Fe is also extremely reactive, producing free radicals when Fe is in excess. For this reason, Fe uptake, storage and allocation has to be properly sensed and tightly regulated to prevent either deficiency or toxicity (Khan *et al*., 2014).

Fe is abundant in most soils, although it is typically found as insoluble Fe^3+^-complexes and therefore unavailable for uptake into root cells (Römheld and Marschner, 1986). In turn, land plants have evolved two strategies to overcome the challenge of solubilizing and importing Fe into roots from the rhizosphere. Strategy I is a reduction strategy, carried out at the plasma membrane where membrane bound reductases directly reduce Fe^3+^ to Fe^2+^ prior to transport across the membrane by transporters of the ZIP family (Hindt and Guerinot, 2012). Strategy II is a chelation strategy, which is mediated by the release phytosiderophores such as deoxymugineic acid (DMA) into the rhizosphere where a soluble Fe^3+^-DMA complex is formed and imported into roots by transporters of the *Yellow Stripe* family (Curie *et al*., 2001, 2009; Inoue *et al*., 2009; Lee *et al*., 2009).

Dicots, such as *Arabidopsis*, utilize Strategy I and form the core of our understanding of Fe uptake and its regulation during changes in Fe availability. In these plants, Fe^3+^ is initially released from negatively charged soil particles by acidification of the rhizosphere through the P-type ATPase AHA2 (Santi and Schmidt, 2009). Once released from the soil particles, Fe^3+^ is reduced to Fe^2+^ by the transmembrane protein <u>F</u>erric <u>R</u>eduction <u>O</u>xidase (FRO2) (Robinson *et al*., 1999). Finally, Fe^2+^ is transported into the root primarily by the <u>I</u>ron <u>R</u>egulated <u>T</u>ransporter <u>1</u> (IRT1). While IRT1 has high affinity towards Fe^2+^, it is also able to transport a broad range of divalent metals including zinc, manganese, and the non-essential element cadmium (Cd) (Korshunova *et al*., 1999).

Transcriptional regulation of the Fe uptake machinery is primarily mediated by the *FIT network*, which consists of five bHLH transcription factors (TFs) of the subgroup Ib: FIT (bHLH029), bHLH38, bHLH39, bHLH100, and bHLH101 (Yuan *et al*., 2008; Sivitz *et al*., 2012; Wang *et al*., 2013). These genes are under independent regulatory schemes but they are thought to function as homo- and heterodimers, allowing for multiple input signals to induce or repress Fe uptake. During prolonged Fe deficiency, or H_2_O_2_ exposure, the zinc finger transcription factor ZAT12 is induced and represses FIT activity, thereby inhibiting Fe uptake and preventing further Fe excess-mediated oxidative damage (Le *et al*., 2016). Additionally, Fe uptake is regulated by the plant hormones ethylene, cytokinins, jasmonates, auxin, and the signaling molecule nitric oxide (Hindt and Guerinot, 2012). FIT has been shown to interact with additional TFs like EIN1 and EIL3 to promote Fe uptake by stabilizing FIT (Lingam *et al*., 2011; Yang *et al*., 2014) while the presence of nitric oxide prevents the 26S-mediated degradation of FIT (García *et al*., 2010).

While the transcriptional regulation of *IRT1* and associated genes involved in root Fe uptake is mediated by the *FIT network*, a second clade of bHLH proteins, the *PYE network*, regulates intracellular Fe homeostasis. The founding member of this clade, <u>P</u>OPE<u>YE</u> (PYE), was found to be induced by Fe deficiency in roots (Long *et al*., 2010). In addition to PYE itself, the *PYE network* is composed of PYE-like proteins (PYEL) bHLH34, bHLH104, bHLH115, and ILR3 (Zhang *et al*., 2015; Li *et al*., 2016; Liang *et al*., 2017), as well as BRUTUS (BTS), an E3 ubiquitin ligase that degrades PYEL proteins to prevent an excess Fe accumulation (Long *et al*., 2010; Selote *et al*., 2015; Hindt *et al*., 2017). The *PYE network*, like the *FIT network*, is thought to operate by hetero- and homo-dimerization of TFs, which then bind to the promoters of key Fe homeostasis genes such as *ZIF1, NAS4*, and *FRO3* (Long *et al*., 2010), as well as *bHLH38/39/100/101* (Zhang *et al*., 2010; Li *et al*., 2016; Liang *et al*., 2017).

Though the components of Fe uptake machinery in roots have been known for years, the mechanisms behind Fe sensing and signaling at the whole plant level is an ongoing active area of research. *Arabidopsis* is known to have at least two distinct Fe sensing systems, a local sensing system in roots, and a systemic sensing system in leaves (Khan *et al*., 2018). The systemic sensing system allows the leaves to dictate the amount of Fe to be acquired by roots, while the local sensing system allows individual roots to regulate their response to their local environment, mostly through post-translational mechanisms (Barberon *et al*., 2011; Sivitz *et al*., 2011; Guillaume *et al*., 2018).

Cadmium (Cd), on the other hand, is a non-essential element that shares chemical properties with Fe and therefore is capable of entering root cells using the Fe uptake system (Meda *et al*., 2007; Wu *et al*., 2012; Lešková *et al*., 2017). Cadmium has been shown to induce genes such as *IRT1* and *OPT3*, which are usually induced under Fe-limiting conditions; however, whether this Fe deficiency-like response is only due to direct competition and reduced Fe uptake in the presence of Cd, or whether Cd directly impairs the Fe sensing mechanism is currently not known. Recently, there have been significant advances at defining Fe responsive gene networks in a tissue and cell-specific manner (Khan *et al*., 2018). In addition, *Arabidopsis* mutants that constitutively over accumulate Fe in leaves and roots even in the presence of Cd have also been identified (*opt3-2 and opt3-3*). In this work, we used low levels of Cd to probe whether this non-essential element directly impairs Fe sensing in wildtype plants and in a mutant that over accumulates Fe in leaves and roots (*opt3-2*). Our results show that Cd does induce a transcriptional program consistent with Fe deficiency in wildtype leaves and roots. However, many of these genes were not induced by Cd in plants that constitutively over accumulate Fe (*opt3-2*). Notably, despite the presence of high levels of Fe in *opt3* leaves, Cd consistently induced a specific core of Fe responsive genes known to be localized in the leaf vasculature. Further analyses demonstrate that genes originally induced by Cd in wildtype plants, but not in *opt3-2* (Fe excess conditions), belong to networks associated with pathogen responses and oxidative stress. Taken together, our results suggest that when plants experience opposite cues (Fe-deficiency and high H_2_O_2_), there is a *hierarchical regulation* of Fe homeostasis in which H_2_O_2_ overrides the induction of a subset of genes that otherwise would have been induced by Fe-deficiency.

## Materials and Methods

### Plant growth

*Arabidopsis* wildtype Columbia (Col-0) and *opt3-2* plants were germinated on ¼ MS agar plates after two days of stratification at 4°C in the dark. After approximately 10 days plants were transferred to replete hydroponic media as previously described (Khan *et al*., 2018). Solution was changed every two days and aerated with small aquarium type air pumps. At bolting, fresh solution with indicated concentrations of CdCl_2_ was added for 72 hrs.

### RNA sequencing and data analysis

Leaves and roots were harvested separately, pulverized in a mortar and pestle cooled with liquid nitrogen. mRNA was purified using a EZ Plant RNA kit (Qiagen/Germany) and contaminant DNA was removed using a TURBO DNase kit (Invitrogen/USA). Total RNA was submitted to the University of Missouri Core Facility for 100bp Illumina sequencing. The resulting reads were trimmed such that all bases were called at a 95% accuracy using ShortRead (Morgan *et al*., 2009) and mapped to the TAIR10 genome release using Tophat (Kim Daehwan *et al*., 2013). The remaining analysis was carried out in R and Bioconductor (Huber *et al*., 2015; R Core Team, 2018). Feature counting was performed using ShortRead (Morgan *et al*., 2009) and 92% of raw reads were uniquely mapped. Differential expression was called using edgeR (Robinson *et al*., 2009), heatmaps and Venn diagrams were generated in gplots (Warnes *et al*., 2016). Ontology enrichment tests were performed using GOstats (Falcon and Gentleman, 2007) using the conditional hypergeometric test with a p-value cutoff of 0.05. The union of the Fe excess dependent and Fe excess independent was used as the gene universe for each test, for leaves and roots independently. Previously published sequencing data (without Cd treatment) can be found on NCBI GEO GSE79275, while Cd exposed sample can be found as GSE128156. These datasets comprise the SuperSeries GSE128157.

### Assembly of the *expanded root ferrome* and *leaf Ferrome*

The *expanded root ferrome* and *leaf ferrome* (Table S1) were assembled using tissue specific Fe deficiency microarray profiling experiments from Stein and Waters, 2012 (ecotypes Kas and Tsu after 24 and 48hr Fe deficiency), and Kumar *et al*., 2017 (ecotype Col-0 after 72hr Fe deficiency). Each dataset was analyzed using GEO2R (Barrett *et al*., 2013), fold changes associated with Benjamini and Hochberg moderated p-values (Benjamini and Hochberg, 1995) greater than 0.05 were set to zero. For the Kumar dataset, the expected direction of regulation was determined by the sign (positive or negative) of the log_2_ fold change. For the Stein dataset, an expected direction of regulation was determined for each ecotype by taking the sign of the latest non-zero fold change. A consensus direction of regulation was determined by first constructing an array of fold change signs for each ecotype and sorted under the following rules: genes with two zero-fold changes are declared no-change, one zero fold change and two similarly signed fold changes are declared up or down according to the similar fold changes, genes with no zero fold changes are declared up or down according to the most prevalent sign. The previously published *ferrome* was appended to genes identified in roots. The direction of regulation of *ferrome* genes not identified here were inferred from (Buckhout *et al*., 2009).

### Hydrogen peroxide quantification

Hydrogen peroxide (H_2_O_2_) concentrations in leaves were determined by the method described in (Velikova *et al*., 2000). Briefly 500mg fresh tissue was homogenized on ice in a solution of 0.1% tri-chloroacetic acid and debris pelleted at 12,000xg. 0.5mL supernatant was added to 0.5mL of 10mM potassium phosphate buffer (pH 7) and 1mL 1M KI. The absorbance at 390nm and compared to a standard curve of H_2_O_2_. In roots H_2_O_2_ was quantified using the Amplex Red reagent (Thermo Fisher Scientific, USA), according to the methods of Brumbarova *et al*.(2016). Oxidation of Amplex Red to resorufin was measured by quantifying resorufin fluorescence with excitation at 545nm and emission at 590 nm.

### Photosynthetic activity

Col-0 and *opt3-2* were stratified at 4°C in the dark and subsequently germinated on ¼ MS media plates under long-day conditions (16h light/8h dark). After 14 days of growth, seedlings were transferred either to fresh ¼ MS plates (mock control) or to ¼ MS plates supplemented with 20 μM Cd. Subsequently, we started to monitor plant photosynthetic performance on a daily basis by determining maximum quantum yield of photosystem II (*Fv/Fm*) using a Walz Imaging PAM (Kunz *et al*., 2009).

### Purification and spectroscopy of AtNEET

AtNEET stability assays were conducted at room temperature (pH 8.0) and the indicated Cd concentrations with 100 μM of purified AtNEET as previously described (Nechushtai *et al*., 2012). Briefly, AtNEET protein was expressed in *E.coli* BL21 competent cells. At an OD_600nm_ of 0.6, 750 μM FeCl_3_ was added to the cell culture and incubated for an additional 16 h at 23°C. AtNEET was purified from lysed cells using Ni-NTA column and size exclusion chromatography, as described by Nechushtai *et al*. (2012). UV-Vis absorption spectroscopy was performed on a Cary50 spectrometer equipped with a temperature controlled cell.

## Results

### Cadmium induces a transcriptional Fe deficiency-like response in roots and leaves

In this work, we took advantage of the shared chemical properties of Fe and Cd, and use Cd as a *chemical probe* to assess Fe deficiency responses. Since plant responses to cadmium vary depending on its concentration, we began this work by identifying a Cd concentration where the visual damage to leaves (i.e. chlorosis) was minimal. We included in these experiments the *Arabidopsis* mutant *opt3-2*, which constitutively over-accumulates Fe in leaves even in the presence of Cd (Mendoza-Cózatl *et al*., 2014; Zhai *et al*., 2014), thus offering a suitable background to explore Cd-induced Fe deficiency-like responses in the presence of high levels of Fe. Wildtype and *opt3-2* plants were grown in replete hydroponic solution to bolting stage (approx. 4 weeks) and then exposed to several concentrations of CdCl_2_ for 72 hr (Fig. 1). While high concentrations of Cd (> 50 μM) induced leaf yellowing and necrotic lesions, exposure to up to 20 μM CdCl_2_ for 72 hrs had minimal impact on plant morphology in both genotypes, wildtype and *opt3-2*. Therefore, we selected 20 μM CdCl_2_ for further experiments.

**Fig. 1:**
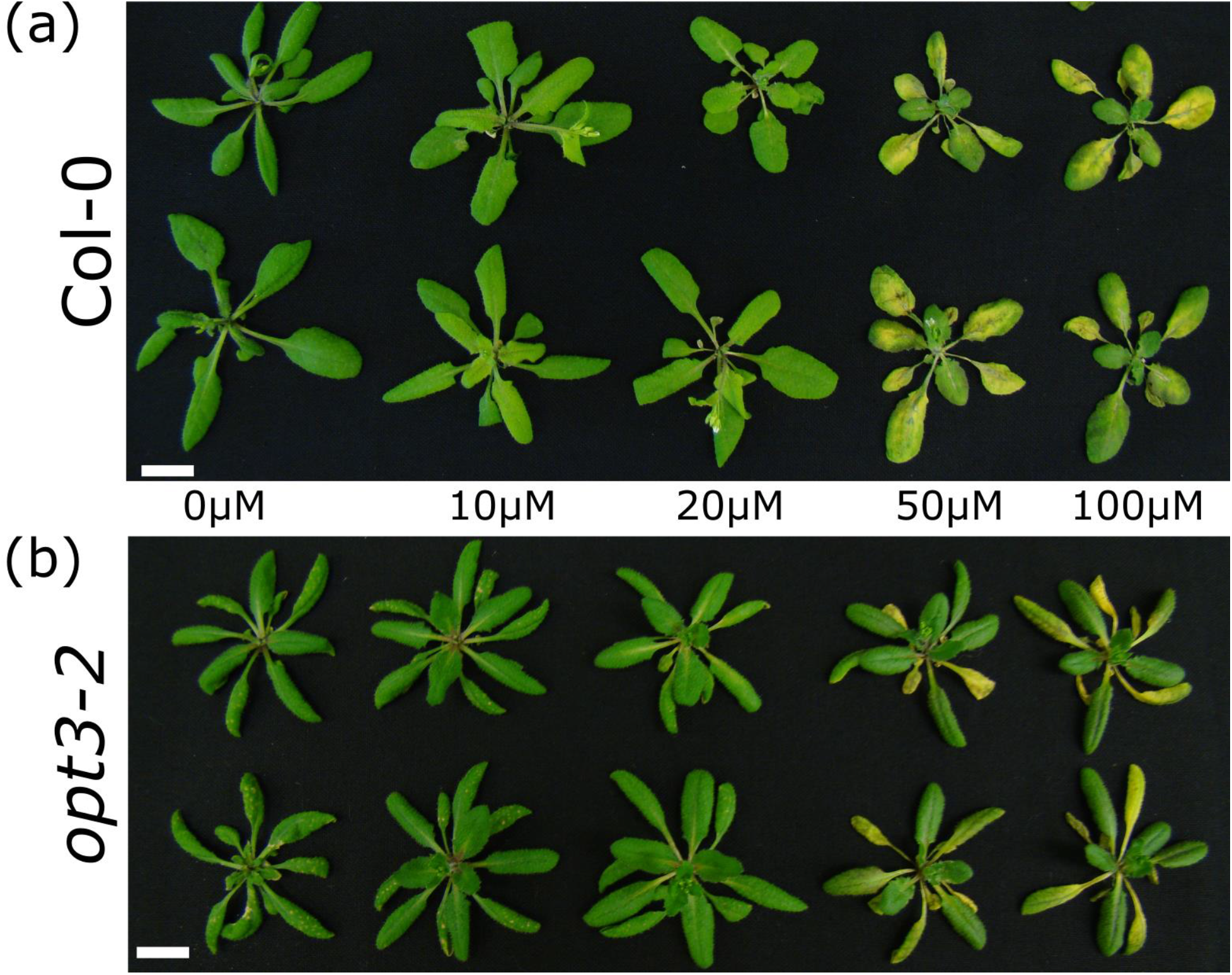
Visual phenotypes of wildtype and *opt3-2* Arabidopsis plants exposed to cadmium for 72 hrs. **(a)** Col-0 (wildtype) and **(b)** *opt3-2* plants were grown in replete media for 4 weeks before being exposed to different Cd concentrations for 72 hrs. 20 μM Cd was the highest Cd concentration at which visual toxicity symptoms were minimal in both genotypes.

To begin dissecting the wildtype and *opt3-2* responses to Cd exposure, within the context of Fe deficiency, we conducted whole-genome transcriptome analyses of leaves and roots separately. Three biological replicates of each tissue/genotype were used for Illumina sequencing and after removal of low confidence base pairs and short reads, 716 million reads were used for calling differential expression under the over dispersed binomial model implemented in edgeR (Robinson *et al*., 2009). To minimize unreliable fold changes, only genes with at least 50 reads in at least one condition and absolute log_2_ fold changes ≥ 0.5 were considered for statistical analyses (Table S2). In wildtype plants, Cd induced changes in 3,292 genes in leaves (46% induced) and 4,256 genes in roots (49% induced) (Fig. S1). A similar number of genes (3,735; 46% induced) were differentially expressed in *opt3-2* leaves; however, Cd induced a substantial deregulation of transcripts in *opt3-2* roots, totaling 6,527 differentially expressed genes, of which 42% were induced (Fig. S1). Gene expression profiles and comparisons between genotypes can be visualized in absolute (FPKM) or relative (log_2_ fold changes) mode through a stand-alone version of an electronic Fluorescent Pictograph Browser available at http://gene.rnet.missouri.edu/efp/cgi-bin/public_html/efpWeb.cgi

To determine the extent of the Fe deficiency response induced by Cd, we compared the identity of Cd deregulated genes against datasets specific to leaves and roots containing genes deregulated (i.e. induced or repressed) under true Fe deficiency conditions. These datasets include an extended version of the published *ferrome* for roots (Schmidt and Buckhout, 2011) and a leaf-specific data set that contains genes consistently affected by Fe availability across several transcriptome datasets where leaves were analyzed separately from roots (Stein and Waters, 2012; Kumar *et al*., 2017). In total, the leaf dataset (*leaf ferrome*, Table S1) included 239 genes (172 induced, 67 repressed) while the root dataset (Table S1) contains 406 genes (250 induced, 156 repressed). By using these datasets, we were able to assess the degree of Fe deficiency response elicited by Cd. For instance, in wildtype leaves we found a significant overlap between Fe deficiency responses and Cd exposure, with 82 genes induced and 41 repressed. These numbers represent 48% (induced genes) and 61% (repressed genes) of the true Fe deficiency response from the *leaf ferrome* (Fig. 2a). Examples of genes deregulated by Cd and Fe in leaves include *IMA3* (upregulated ~5 log_2_ fold), which encodes a short polypeptide known to be involved in Fe deficiency responses (Grillet *et al*., 2018; Hirayama *et al*., 2018), the jasmonic acid signaling marker *PDF1.2* (induced ~9 log_2_ fold) (Ahmad *et al*., 2011; Zarei *et al*., 2011; Cabot *et al*., 2013), and *FER4* (repressed ~2 log_2_ fold), which encodes a ferritin isoform. Not all Fe deficiency related transcripts were found to be deregulated by Cd exposure. For example, neither *CGLD27* (Urzica *et al*., 2012), or a key regulator of salicylic acid response *SARD1* were induced (Wang *et al*., 2011). Similar trends were found in roots, 118 genes were induced and 60 repressed, which represent 47% of induced and 38% of repressed genes present in the *expanded root ferrome* (Fig. 2b). These results suggest that Cd induces a partial but significant Fe deficiency-like response in both leaves and roots.

**Fig. 2:**
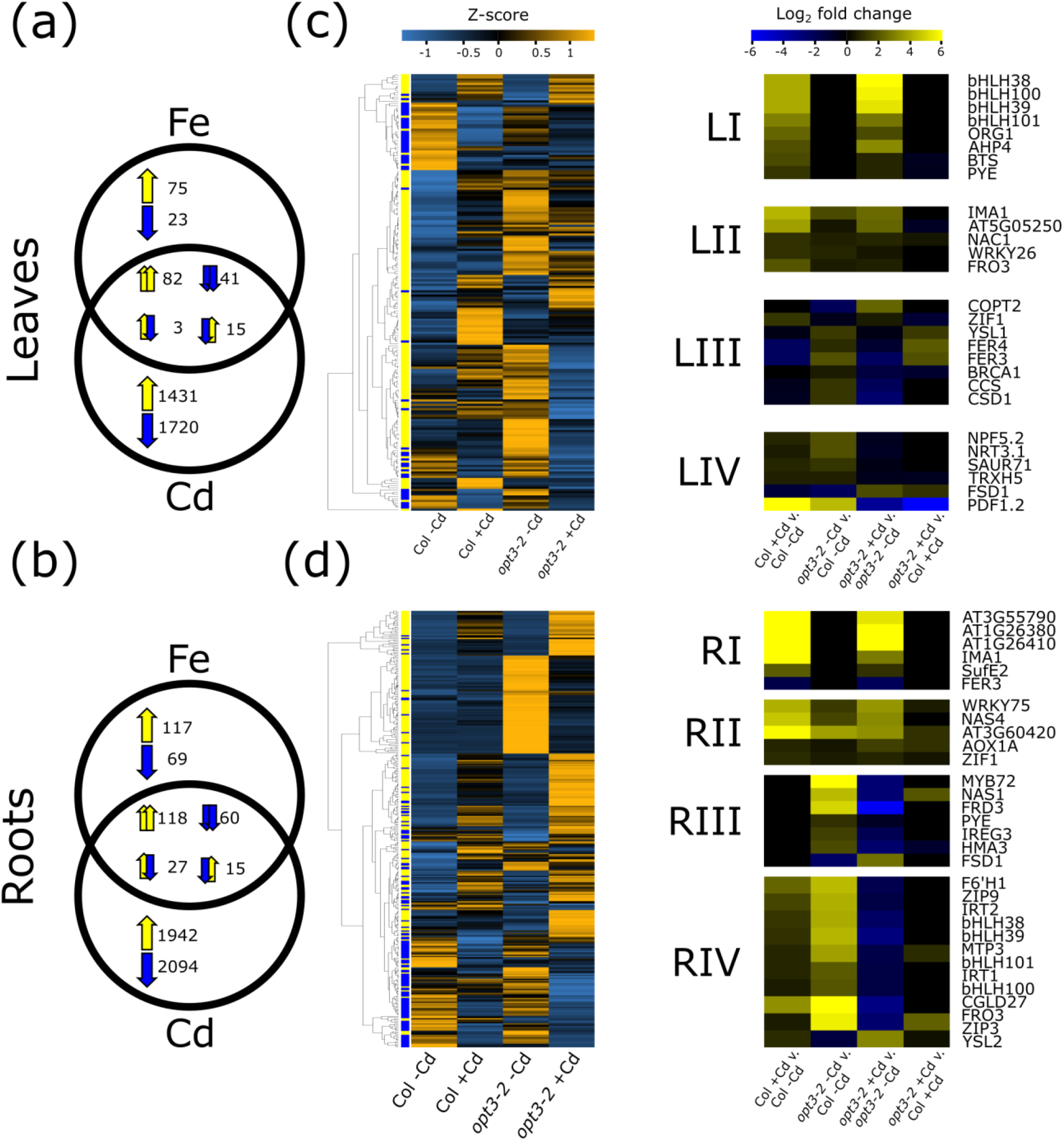
Cadmium induces an iron deficiency response in leaves and roots. **(a-b)** Of all of the genes that are consistently responsive to Fe limitation in wildtype Arabidopsis plants, approximately half of them are also responsive to Cd in leaves **(a)** and roots **(b)**. The pair of arrows in each intersection indicate induction/repression for Cd exposure (left arrow within each pair) and Fe deficiency (right arrow within each pair). **(c-d)** Genes deregulated by Fe deficiency and Cd exposure in wildtype plants and *opt3-2* were hierarchically clustered according to their log_2_ fold changes and colored according to their gene-wise Z-scores for leaves **(c)** and roots **(d)**, left panels. Clusters with specific trends were grouped into four groups for leaves and roots (right panels, LI-RV) and shown as Log_2_ fold changes between genotypes and treatments.

### Fe overload partially restricts the Cd-induced Fe deficiency response

Cadmium-induced Fe-deficiency responses are often attributed to the competition of Fe and Cd for the same root uptake system [i.e. IRT1 (Connolly *et al*., 2002; Lešková *et al*., 2017)]. Therefore, we decided to explore if Cd was still able to induce an Fe deficiency-like response in tissues with high levels of Fe. The *Arabidopsis* mutant *opt3-2* has been shown to over accumulate Fe in leaves and roots in the presence of Cd (Mendoza-Cózatl *et al*., 2014). Moreover, the *opt3-2* leaf transcriptome is consistent with an adequate sensing of Fe excess, while roots display a constitutive Fe deficiency response despite accumulating high levels of Fe (Khan *et al*., 2018). Therefore, we employed hierarchical clustering of all genes differentially expressed in at least one comparison between the four treatment groups: wildtype and *opt3-2* plants exposed or not to 20 μM CdCl_2_ (Fig. 2c, d).

To streamline the simultaneous inspection across all genotypes and treatments, the clustering scheme is presented as the mean of <u>c</u>ounts <u>p</u>er <u>m</u>illion (CPM). Using this approach, we were able to identify several distinct patterns for leaves [LI to LIV] and roots [RI to RIV] (Fig. 2c, d; Table S2). Briefly, Group I consists of genes which are similarly affected by Cd in both wildtype and *opt3-2*. Genes in group II are deregulated in *opt3-2* relative to Col-0 prior to Cd treatment but they are similarly induced after Cd exposure. Group III comprises genes which were initially deregulated in *opt3-2* relative to wildtype in the absence of Cd but interestingly, Cd exposure reversed the sign of these genes in *opt3-2*. Finally, genes in Group IV have a similar pattern as Group III but the magnitude of the changes was substantially different. Specific examples of genes within these clusters can be further separated in a tissue-specific manner. For instance, Group LI in leaves contains vascular localized genes such as *bHLH38, bHLH39, bHLH100*, and *bHLH101* as well as *PYE, BTS*, and *ORG1*. All these genes were induced by Cd independent of the background genotype (Fig. 2c). *FRO3* however, is induced by Cd exposure in wildtype and *opt3-2* but it was already induced in *opt3-2* prior to Cd exposure hence its placement in group LII. Group LIII represents a very interesting group of genes including *CSD1* and its chaperone *CCS, YSL1, FER3/4*, and *ZIF1*. These genes were induced in *opt3-2*, but when Cd stress was imposed, they switched from induction to repression. Group LIV contains genes such as PDF1.2, an ethylene and jasmonate responsive plant defensin which is highly induced by Cd stress, induced in the *opt3-2* background, but repressed once *opt3-2* is treated with Cd.

The transcriptional program elicited by Cd in roots was far more complex in both genotypes than the leaf patterns. For instance, gene group RI did not contain the Ib bHLHs as found in LI (Fig. 2c, d), instead the transmembrane protein AT3G55790 and two FAD-binding Berberine family proteins (AT1G26380 and AT1G26410) were found to be highly induced, as well as sulfur E2 (SufE2), a protein likely to play a role in FeS cluster assembly (Narayana *et al*., 2007). Group RII contains genes such as *NAS4* and the vacuolar Zn-nicotianamine importer *ZIF1*, both of them induced by Cd, indicating that the plant is activating a heavy metal sequestration program. *AOXA1*, which is a component of the alternative oxidase branch of mitochondrial respiration, was also found in group RII. AOX expression is correlated with oxidative stress and H_2_O_2_ concentrations and is expected to be involved in retrograde stress signaling from the mitochondria to the nucleus (Saha et al., 2016). Group RIII contains several genes of interest for Fe homeostasis, most notably the transcription factors *PYE* and *MYB72*, which are differentially regulated by Cd in the roots of *opt3-2* mutants but not of wild type plants. In *opt3-2* roots, these genes are constitutively highly expressed in the absence of Cd, but are repressed to wildtype levels in the presence of Cd (Fig. 2d). Other Fe related allocation genes such as *IREG3, HMA3*, and *FRD3* are also included in this RIII group. Finally, RIV contains the *Fe regulon* including *bHLH38, bHLH39, bHLH100, bHLH101, IRT1*, and *F6’H1* as well as the transporters *ZIP3, ZIP9*, and *MTP3*. These genes are all induced by Cd exposure, strongly induced in *opt3-2* prior to Cd exposure but repressed after Cd exposure. These and additional transcriptional responses of key genes were further validated by qRT-PCR (Fig. S2, Table S5). Altogether, these results suggest that Cd does triggers a Fe deficiency-like response in wild type plants, at mild concentrations; however, when combined with other stresses such as Fe excess, different or even opposite transcriptional programs are activated. Notably, this *hierarchical* regulation of Fe homeostasis seems to apply only to a very specific set of genes and by imposing different levels of stress, Cd and Cd + Fe excess (i.e. *opt3-2*), we were able to separate clusters with distinct transcriptional patterns.

### Fe deficiency responses are hierarchically regulated based on multiple inputs

In the presence of Fe excess, Cd elicits different responses of Fe deficiency markers in leaves and roots (Fig. 2c, d). While the Cd induction of the subgroup Ib *bHLHs* even in the presence of high levels of Fe (i.e. *opt3* leaves) indicates that Cd interferes with Fe sensing, the repression of the *FIT network* by Cd in *opt3-2* roots indicates that the Fe deficiency signals can be overridden by other mechanisms (see Group RIV in Fig. 2d). In order to better separate the behavior of these Cd-induced Fe-deficiency responsive genes, we clustered the *ferrome* genes according to their regulation in response to Cd for each genotype (i.e. Col or *opt3-2*). Genes which are similarly induced in each genotype were classified as Fe excess *independent* (Fig. 3, orange dots), as they were induced by Cd despite the high Fe levels in *opt3-2* (Groups I and II), while those whose regulatory pattern did change (Groups III and IV) were classified as Fe *excess-dependent* (Fig. 3, purple dots). Both leaves and roots displayed a combination of Fe *excess-dependent* and Fe *excess-independent* gene expression (Fig. 3 a, b). Interestingly, some genes whose transcriptional response to Cd was found to be Fe excess-independent in one tissue, were found to be Fe excess-dependent in the other. Notably, genes of the subgroup Ib bHLHs were found to be Cd-inducible and Fe *excess-independent* in leaves, whereas the same group displayed a different expression pattern in roots: they were induced by Cd in wildtype root systems but repressed in *opt3-2* (i.e. in *opt3-2* + Cd; Fig. 3b).

**Fig. 3:**
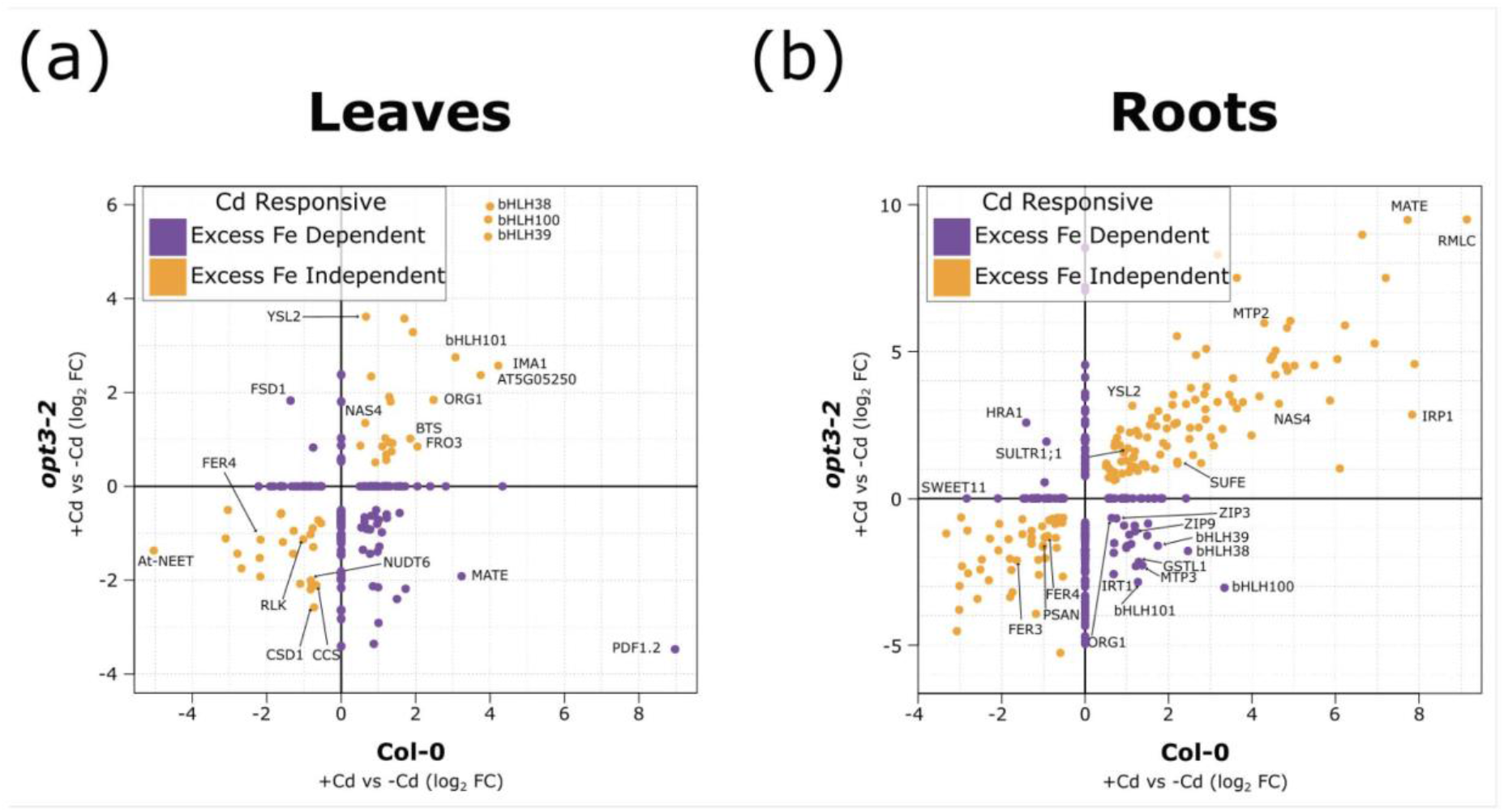
Cadmium responsive genes are differentially regulated in a tissue-specific manner depending on the Fe levels in plant tissues. **(a)** Leaf and **(b)** root Fe deficiency markers similarly induced/repressed by Cd in wildtype and *opt3-2* are labeled as “Fe excess independent” and shown as orange dots. Fe deficiency markers with opposite expression patterns in wildtype and *opt3-2* during Cd exposure are labeled as “Fe excess dependent” and shown as purple dots.

To further uncover a possible signaling mechanism responsible for separating the Cd-inducible Fe excess-*dependent* cluster from the Fe *excess-independent* cluster, we used gene ontology (GO) enrichment. This approach allows the identification of broad trends associated to the role of each gene within clusters (Fig. 4). The result from this analysis showed that, in leaves, the Cd-inducible Fe *excess-independent* cluster was enriched in terms related to Fe or metal ion homeostasis (Fig. 4a), while the Cd-inducible Fe excess-*dependent* cluster was significantly enriched in terms related to biotic stress (Fig. 4b). Genes related to biotic stress have frequently been observed in mRNA profiling experiments studying Cd stress - an abiotic stress - but a rationale behind this trend is still lacking. Our results however, suggest that the mechanism separating the Fe excess-*dependent* and *-independent* clusters is closely related to biotic stress responses, which often rely on reactive oxygen species (ROS) for signaling and defense during a pathogen infection. In roots, the Fe *excess-independent* cluster was enriched in terms related to secondary metabolism (Fig. 4c), while the Fe excess-*dependent* cluster in roots was enriched with terms relating to heavy metal homeostasis and, again, biotic stress response terms (Fig. 4d). This trend also suggests that the mechanism repressing Fe responsive genes in leaves and roots under Cd exposure is different. In leaves, this may be the result of Cd interfering with Fe sensing, while in roots, a secondary signaling mechanism likely related to ROS, appears to exert additional transcriptional control on the Fe transcriptional network.

**Fig. 4:**
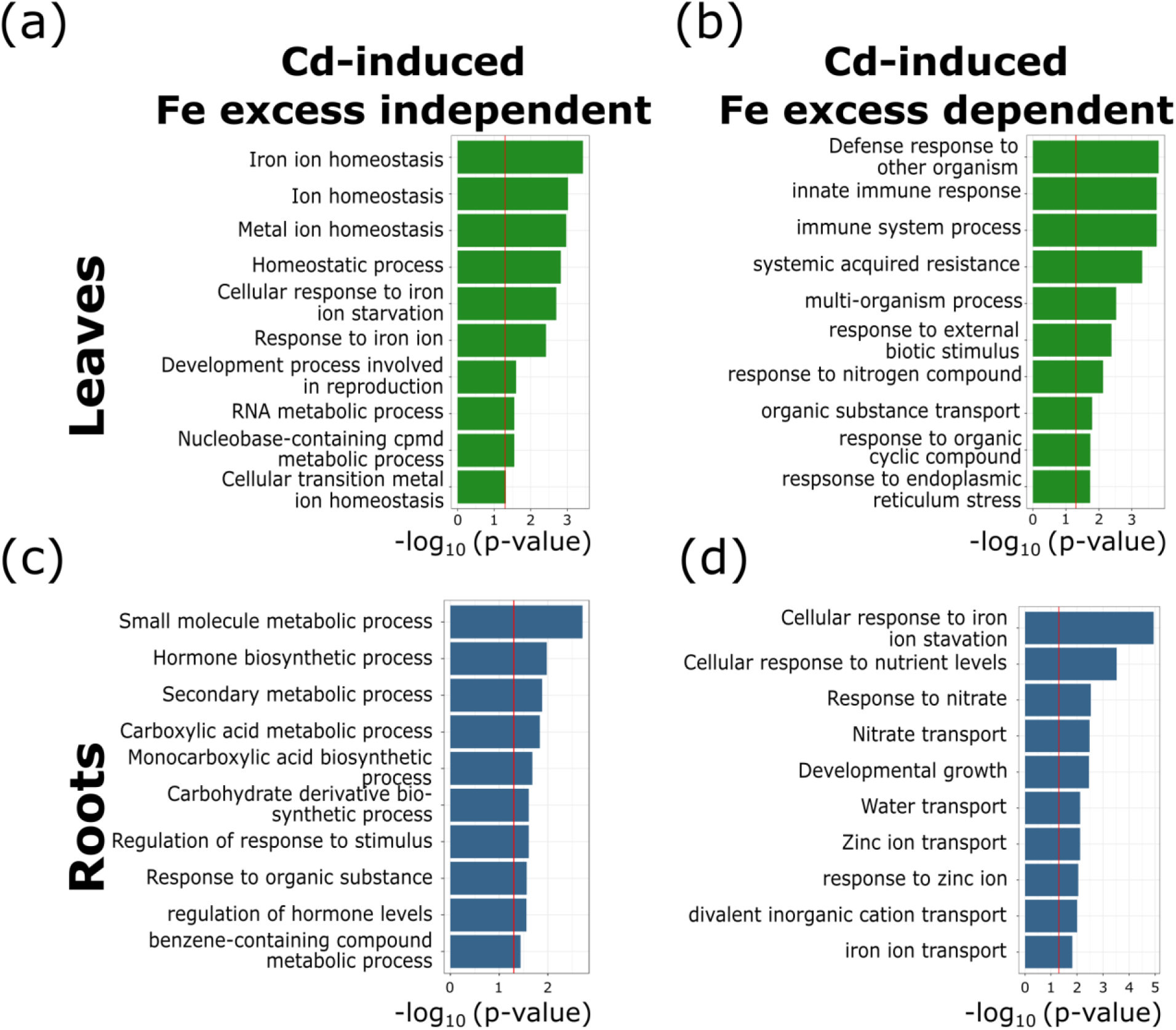
Classification of cadmium-induced genes based on their biological process and sensitivity to Fe excess. Gene ontology enrichment tests were performed for **(a,c)** Cd induced Fe excess independent and **(b,d)** dependent clusters in leaves and roots. Red line indicates significance threshold (p = 0.05).

### Iron excess and cadmium have additive effects on H_2_O_2_ accumulation

Our RNA sequencing data consistently show that Cd elicits an Fe deficiency-like response by upregulating a specific set of transcripts, even in the presence of Fe excess, but also that a significant number of genes originally induced by Cd were repressed to wildtype levels when Fe was in excess (i.e. in *opt3-2*). These repressed transcripts are disproportionately associated with biotic stress responses, and likely share a regulatory component. Reactive oxygen species, such as H_2_O_2_, are generated by the <u>R</u>espiratory <u>B</u>urst <u>O</u>xidase NAD<u>PH</u> protein <u>D</u> (RBOHD) during pathogen attack to mediate the defense response (Miller *et al*., 2009; Pogany *et al*., 2009; Maruta *et al*., 2011; Torres *et al*., 2013). In leaves, *RBOHD* was induced by Cd in wildtype and induced in *opt3-2* without Cd exposure, to levels similar to wildtype exposed to Cd (Fig. S3). Hence, we hypothesized that the repression of some Fe deficiency markers in *opt3-2* after Cd exposure may be the result of a H_2_O_2_-mediated transcriptional reprogramming.

To support this hypothesis, H_2_O_2_ levels were measured in leaves and roots of wildtype and *opt3-2* plants exposed or not to Cd (Fig. 5). In leaves, exposure to 20 μM Cd had no impact on H_2_O_2_ levels, which is which is consistent with our initial observations that at the times tested, 20 μM Cd has no major impact on leaf metabolism (Fig. 1). *opt3-2* plants however, had significantly more H_2_O_2_ compared to wildtype plants and this difference was more pronounced in the presence of Cd (Fig. 5b). The higher levels of ROS in *opt3-2* exposed to Cd, together with the enrichment of ROS-associated genes found through the GO enrichment analysis (Fig. 4a, b) provides a mechanism to explain why some Fe responsive genes initially induced by Cd in wildtype plants were later repressed by Cd in *opt3-2* (i.e. the Cd inducible Fe excess-dependent cluster, Fig. 3a). Furthermore, it also suggests that when plants experience simultaneously Fe deficiency-like conditions and high ROS, there is a *hierarchical regulation* of Fe deficiency responses where ROS prevent the induction of genes that otherwise would have been induced as part of the Fe deficiency response in leaves. Perhaps more interesting is the fact that in leaves, and only in leaves, bHLHs of the subgroup Ib are insensitive to this ROS-mediated *hierarchical* regulation (Fig. 3a). The H_2_O_2_ levels in roots followed the same trend as in leaves, but the magnitude of changes was more dramatic. For instance, Cd exposure elevated the H_2_O_2_ concentration in wildtype roots to similar levels found in unexposed *opt3-2*. In addition, Cd increased the H_2_O_2_ levels in both genotypes but the concentration of H_2_O_2_ in *opt3-2* was significantly higher than wildtype. This higher H_2_O_2_ content in *opt3-2*, induced by Cd, may also provide the basis to explain some of the distinct transcriptome profiles observed in roots, where Cd-exposed wildtype and unexposed *opt3-2* show similar expression patterns (Fig. 2d, cluster RII), but the high levels of H_2_O_2_ in Cd exposed *opt3-2* represses the expression of Fe regulon.

**Fig. 5:**
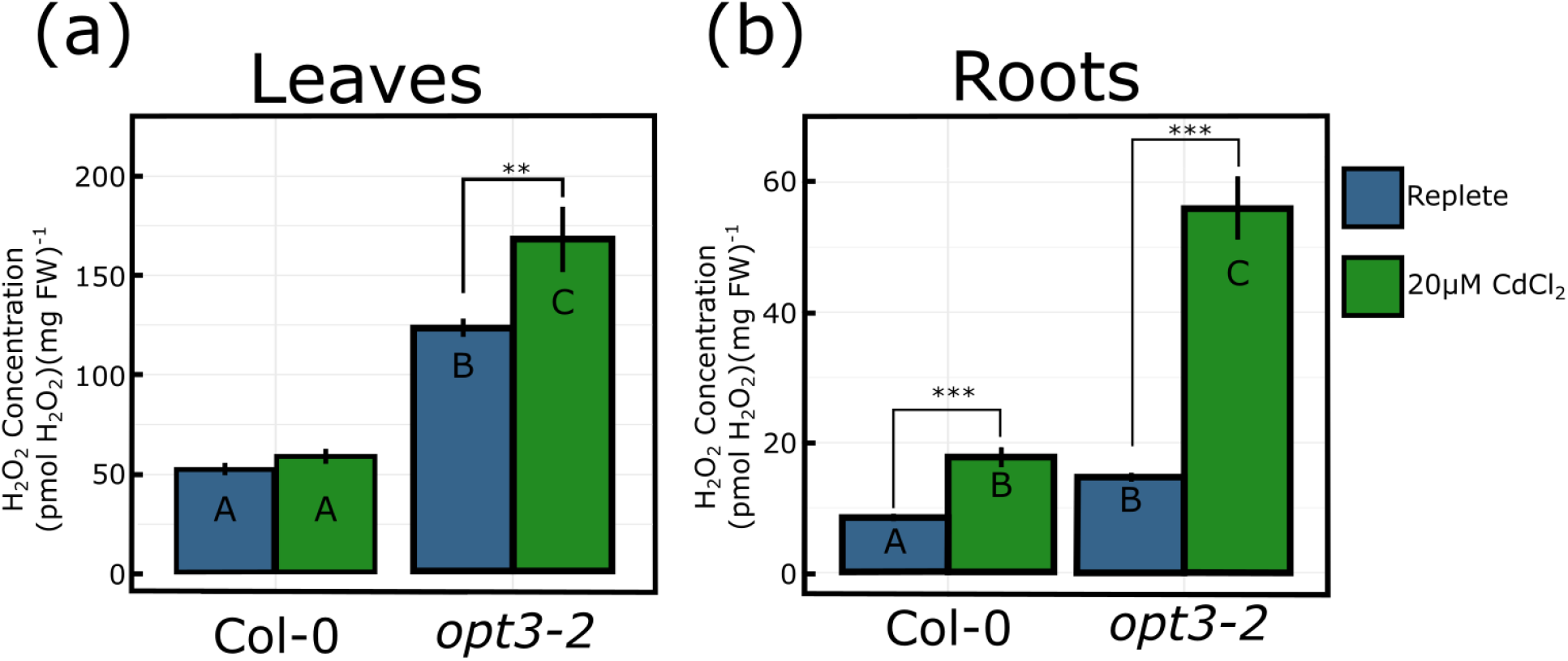
H_2_O_2_ quantification of leaves and roots in wildtype and *opt3-2* plants exposed or not to 20 μM CdCl_2_ for 72 hr (*** indicates p < 0.001, n = 6-10).

### Impaired photosynthetic efficiency and Fe-S metabolism as sources for elevated H_2_O_2_ levels in *opt3-2*

Cadmium is unable to produce reactive oxygen species (ROS) through the Fenton reaction (Strlič *et al*., 2003); however, Cd does generate ROS through displacement of Fe from the active site of proteins and molecules, and photosynthesis is known to be particularly sensitive to Cd toxicity (Küpper *et al*., 2007; Parmar *et al*., 2013). To determine if the increased H_2_O_2_ levels in Cd-exposed *opt3-2* leaves could be explained in part by an impaired photosynthetic apparatus, the maximum potential quantum efficiency of photosystem II (*F_v_/F_m_*) was measured in wildtype and *opt3-2* plants using an Imaging-PAM system (Fig. 6a). The results show that photosynthesis in *opt3-2* remained unaffected under control conditions; however, a 20% reduction in *F_v_/F_m_*, predominantly originating from younger leaves, was found after Cd exposure (Fig. 6b, c).

**Fig. 6:**
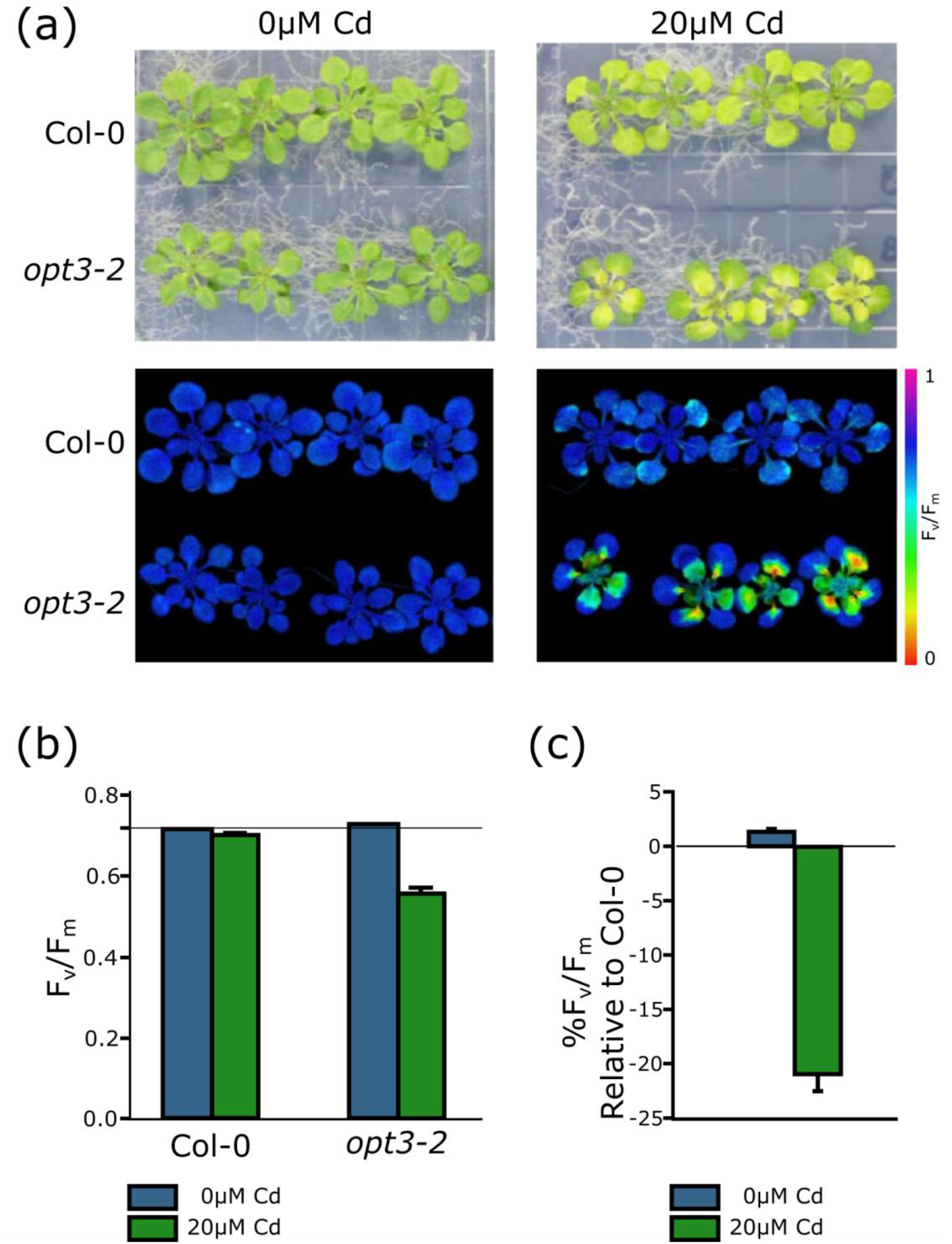
Photosynthesis in *opt3-2* plants is particularly sensitive to cadmium. Photosynthetic efficiency of Col-0 (wildtype) and *opt3-2* was measured in the presence or absence of 20 μM CdCl_2_. **(a)** RGB (top) and *Fv/Fm* false color images (bottom) shows increased chlorosis and photosynthetic inhibition in *opt3-2*. **(b)** Photosynthetic efficiency and **(c)** reduction in photosynthetic efficiency in wildtype and *opt3-2* plants exposed or not to 20 μM CdCl_2_. Error bars indicate standard error of eight individual plants were meaured in each experiment.

Iron-sulfur (Fe-S) metabolism has also been implicated in Fe and ROS homeostasis and more recently AtNEET, an Fe-S donor protein in the chloroplast, has been identified as a key protein regulating Fe-S homeostasis and ROS production. More specifically, the inability of AtNEET to transfer Fe-S clusters triggers an Fe deficiency-like transcript profile (Zandalinas *et al*., 2019). To determine if Cd impairs the stability and/or the ability of AtNEET to transfer Fe-S clusters, we performed AtNEET stability and Fe-S transfer experiments at different Cd concentrations using UV-Vis spectroscopy (Fig. 7). The loss of the characteristic absorption peak of the holo-AtNEET at 458nm shows that AtNEET is a labile protein prone to dissociation in the presence of Cd (Fig. 7a). Next, we asked if Cd could also impair the Fe-S transfer from AtNEET to the acceptor protein ferredoxin (Fd). In the absence of Cd, the transfer of Fe-S clusters between AtNEET and Fd can be tracked by following the 458 to 428 nm spectral shift indicating the Fe-S transfer from the holo-AtNEET to the holo-Fd peak (Fig 7b, inset; (Zandalinas *et al*., 2019)). Notably, the efficiency of this transfer was impaired by Cd (Fig 7b) and quantification of the absorbances at each characteristic peak showed significant decreases in the Fe-S transfer efficiency at Cd concentrations as low as 5 μM Cd (Fig 7c). These results were further confirmed by PAGE analyses where a clear loss of the holo-Fd formation was observed as early as ten minutes after Cd exposure (Fig. S4 a, b). Taken together these data suggest that the elevated H_2_O_2_ levels induced by Cd are the result of an additive impairment of photosynthesis and Fe-S homoeostasis. In turn, the different H_2_O_2_ levels induced by Cd genes in wildtype and *opt3-2* provides a possible mechanism to explain the particular clustering of Cd-induced Fe excess-*dependent* and *-independent* genes observed between wildtype and *opt3-2* genotypes.

**Fig. 7:**
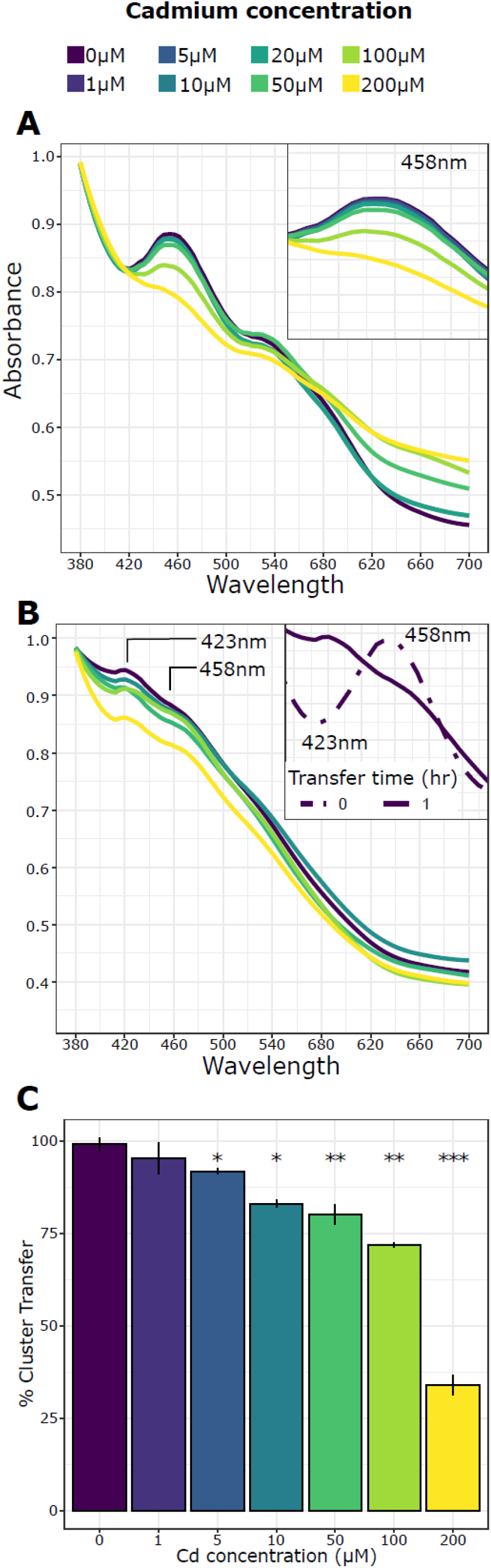
Exposure to Cd impairs AtNEET stability and its capacity to transfer Fe-S clusters to ferrodoxin. **(A)** Absorbance spectra of recombinant AtNEET protein six hours following the addition of Cd, the separation of the peak characteristic absorption at 458nm reflects the loss of the Fe-S cluster from AtNEET. **(B)** In the presence of ferredoxin (characteristic peak 423nm) AtNEET is able to transfer its Fe-S in 1hr (inset) which is impaired by increased Cd concentrations. **(C)** Percent of Fe-S clusters transfered from AtNEET to ferredoxin from (B), * indicates p < 0.01, ** indicates p < 0.001, *** indicates p < 0.0001.

## Discussion

Cadmium has long been known to induce genes that are upregulated when plants experience Fe deficiency and traditionally this has been attributed to a reduced Fe influx into the plant as Cd compete with Fe for the same uptake mechanism in roots (i.e. IRT1). Here we provide evidence suggesting that Cd also interferes with Fe sensing thus leading to transcriptional programs consistent with an Fe deficiency response. Our data also suggest that Fe deficiency responses are regulated by different inputs in a tissue-specific manner, and that some Fe deficiency signals, particularly in roots, can be partially overridden by oxidative stress. This *hierarchical regulation* of Fe homeostasis would prevent further oxidative damage when plants experience opposite cues such as Fe deficiency in the presence of high levels of H_2_O_2_.

### Fe deficiency responses are regulated by several and independent inputs

To test the extent of the Fe deficiency response induced by Cd, we first compiled different datasets and assembled root- and shoot-specific lists of genes consistently deregulated in response to Fe deficiency (*expanded* root *ferrome* and the *leaf ferrome*, Table S1). After comparing these tissue-specific *ferrome* datasets with our Cd-induced transcriptome in wildtype plants, we found a large overlap of genes deregulated by Cd, ~60% of the Fe deficiency markers, including members of the well-known *FIT network, PYE network* and a recently described family of peptides that regulate Fe deficiency responses known as IMAs (Fig. 2, Table S2). If these responses were only the result of uptake competition between Cd and Fe at the root level, this Cd-induced transcriptional response should not occur in plants that accumulate high levels of Fe within their tissues (i.e. *opt3* mutants). However, for some genes, Cd still induced a strong Fe deficiency response regardless of the high levels of Fe present in plant tissues (Fig. 2C, LI). In other cases, the presence of high Fe levels influenced the magnitude of the transcriptional response to Cd, or caused Cd to have an opposite effect on gene expression compared to wildtype plants (Fig. 3, purple dots). This complex and varied pattern of gene expression indicated that the effects of Cd on genes associated with Fe deficiency were due to more than the direct effect of Cd on Fe uptake. Further clustering allowed us to separate and define specific gene clusters and found that some of them are clearly associated with biotic stress and ROS levels. Notably, specific clusters of genes known to be expressed in specific tissues (i.e. vasculature) were induced by Cd regardless of the Fe and ROS levels in plant tissues. This additional information suggested that Fe sensing and Fe deficiency responses are organized in discrete clusters and that the magnitude and direction of transcriptional responses are cluster-specific based on the actual levels of Fe, ROS and the presence of Cd.

### Cadmium impairs Fe sensing

The precise location of Fe sensing in plants is still unclear, but recent data, including this work, suggest that Fe levels may be sensed independently throughout the plant in a tissue and cell-specific manner [Fig. 2, and (Khan *et al*., 2018)]. For instance, leaves of *opt3* mutants (*opt3-2* and *opt3-3*) over accumulate Fe and consequently *opt3* leaves show a transcriptional program consistent with an Fe overload (Khan *et al*., 2018); however, the phloem sap of *opt3* mutants have half of the Fe levels compared to wildtype plants (Zhai *et al*., 2014). Yet, Fe deficiency markers known to be located in the leaf vasculature such as *bHLH38/39/100/101* are not induced in *opt3-2* leaves (Fig. 2c, LI). These results suggest that the high levels of Fe in the leaf apoplast, outside companion cells, may be sufficient to prevent the induction of Fe deficiency genes in companion cells, even if the Fe levels inside companion cells are low (Zhai *et al*., 2014). Alternatively, Fe excess and Fe deficiency may be sensed by independent pathways and when companion cells receive conflicting information (i.e. low levels of Fe in the cytosol but high Fe levels in the apoplast), Fe excess signaling prevails. Interestingly, our data also found that the *bHLH38/39/100/101* cluster is highly induced by Cd in *opt3-2* leaves (Fig. 2c, LI) and this induction is insensitive to the high levels of H_2_O_2_ detected in *opt3-2* leaves (Fig. 5a). Moreover, this induction requires only trace amount of Cd as *opt3* leaves accumulate minimal levels of Cd compared to wildtype (Mendoza-Cózatl *et al*., 2014). Altogether these results suggest that in leaves, and more specifically in companion cells, Cd impairs Fe sensing and that this vascular-specific signaling pathway is insensitive to negative regulators such as Fe excess.

### Fe deficiency is hierarchically regulated by competing nutrient acquisition and oxidative stress signals

Additional evidence for independent tissue-specific Fe sensing mechanisms came from the finding that the transcription of some genes is oppositely regulated in leaves and roots. For instance, while bHLH38/39/100/101 are Cd-induced and Fe excess-independent in leaves, they are Cd-induced but Fe excess-dependent in roots (Fig. 2d). Notably, this severe repression in roots extends to other well-known Fe-deficiency markers such as ORG1 (Kang *et al*., 2003) and FRO3 (Jain *et al*., 2014), suggesting that transcriptional repression by Cd in *opt3-2* roots is under the control of additional inputs. This apparent inconsistency prompted us to explore the nature of these additional inputs. High H_2_O_2_ levels have been previously shown to inhibit Fe deficiency responses (Le *et al*., 2016) and our H_2_O_2_ measurements show that there were significant differences across genotypes, tissues and treatments (Fig. 5a, b). In particular, *opt3* roots exposed to Cd contained 3 times more H_2_O_2_ than wildtype. This dramatic increase in H_2_O_2_ levels induced by Cd provides a mechanistic explanation for the repression of Fe-deficiency markers in *opt3-2* roots, likely mediated by ZAT12 or proteins with similar function (Le *et al*., 2016). In leaves, Cd also induced significantly higher H_2_O_2_ levels in *opt3-2* than wildtype plants and yet some Fe-responsive genes where highly induced despite the high Fe and H_2_O_2_ levels (Fig. 2c, LI cluster) while others were repressed (Fig. 2c, LII cluster). Chloroplast metabolism and photosynthesis are highly dependent on redox reactions and therefore are highly sensitive to Cd (Parmar *et al*., 2013). In turn, our photosynthetic efficiency measurements confirmed that Cd has a higher inhibitory effect in *opt3-2* leaves compared to wildtype plants (Fig. 6) and suggest that in *opt3-2* leaves, chloroplasts are a significant source of H_2_O_2_. Notably, this increase in ROS was similar to what was observed in the dominant negative H89C AtNEET mutant, which exhibits both higher H_2_O_2_ levels and impaired Fe homeostasis due to the impaired transfer Fe-S clusters from AtNEET to acceptor proteins (Zandalinas *et al*., 2019). This same inhibitory Fe-S transfer effect was found *in vitro* in the presence of low levels of Cd (Fig 7b, c). Hence a Cd-dependent impairment of Fe-S homeostasis, combined with inhibition of photosystem II, provides a mechanistic basis for the elevated levels of H_2_O_2_ and the consequent partitioning of Cd-induced Fe excess-dependent and -independent gene clusters. In summary, our results suggest that when plants experience opposite cues (i.e. Fe-deficiency and high ROS levels), there is a *hierarchical regulation* of Fe homeostasis in which ROS override the induction of specific transcriptional programs that otherwise would have been induced by Fe-deficiency.

## Conclusion

Here, we have shown that low levels of Cd induce a Fe deficiency-like response that goes beyond the classic model of Fe/Cd uptake competition at the root level. We also found that Cd impairs Fe sensing, specifically in the leaf vasculature thus affecting the expression of a distinct vascular gene clusters. Interestingly, we also found that in roots, the *FIT network* and Fe uptake system are repressed by Cd in *opt3-2*, which has previously been shown to have a constitutive Fe deficiency response. This repression was concomitant with a drastic increase in H_2_O_2_ concentrations, suggesting that ROS levels can override the induction of some, but not all, of the genes that mediate Fe uptake. Altogether, our results suggest that Fe deficiency responses in *Arabidopsis* are regulated at multiple levels and by different inputs. Moreover, when plants experience opposite inputs, the level of ROS is critical to define the transcriptional outcome of a large set of Fe responsive genes. Our data also suggest that in leaves, the subgroup Ib bHLH transcription factors is insensitive to this *hierarchical regulation* and belong to a gene cluster closely associated with a leaf Fe sensing core and that this leaf-specific Fe sensing core is particularly sensitive, and becomes impaired, in the presence of trace levels of cadmium.

## Supplementary data

**Fig. S1:**
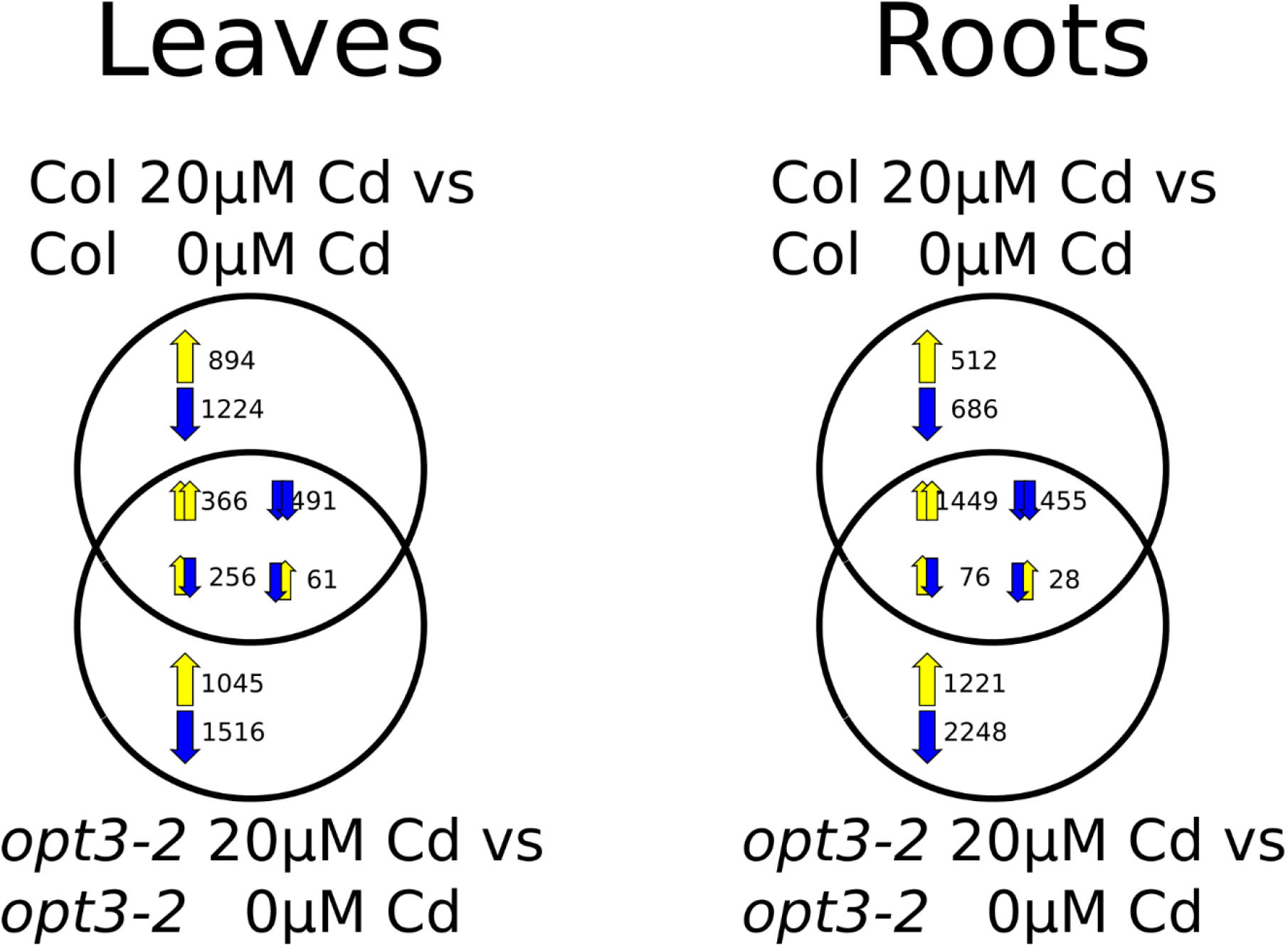
Tissue specific Venn diagrams showing the relationship between Cd exposure in wildtype (Col) and *opt3-2* mutants. Arrows in the intersection represent the direction of regulation for Col (left arrow) and *opt3-2* (right arrow)

**Fig. S2:**
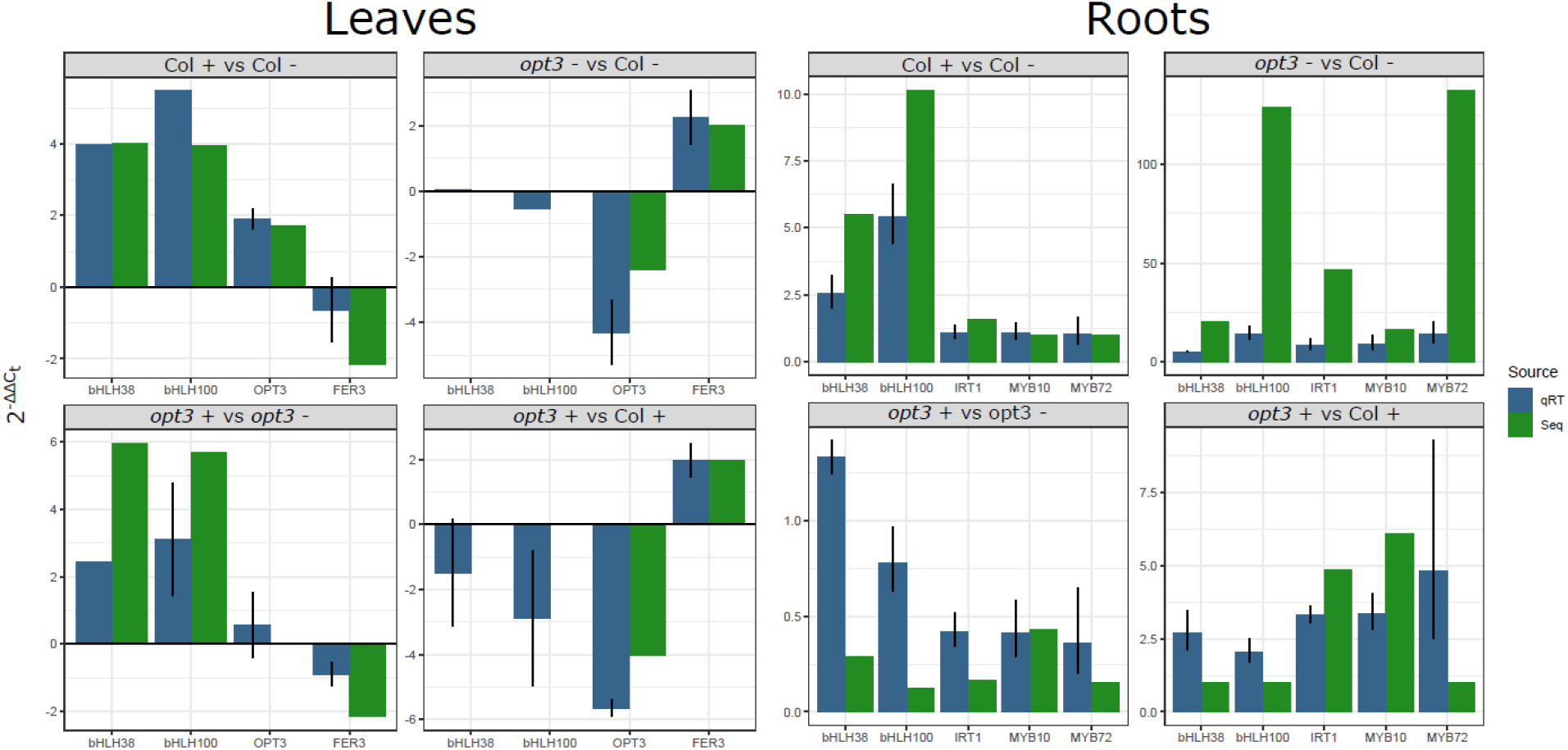
qRT validation of sequencing results. Each gene normalized against ACT 2 expression. Fold changes calculated as 2^−ΔΔ *C_t_*^, and error on the fold change is shown using the combined sample error 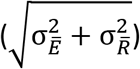 where 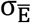 and 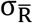 are the sample standard errors, n= 2 with 4-7 plants per replicate. Primers used can be found in Table S5.

**Fig. S3:**
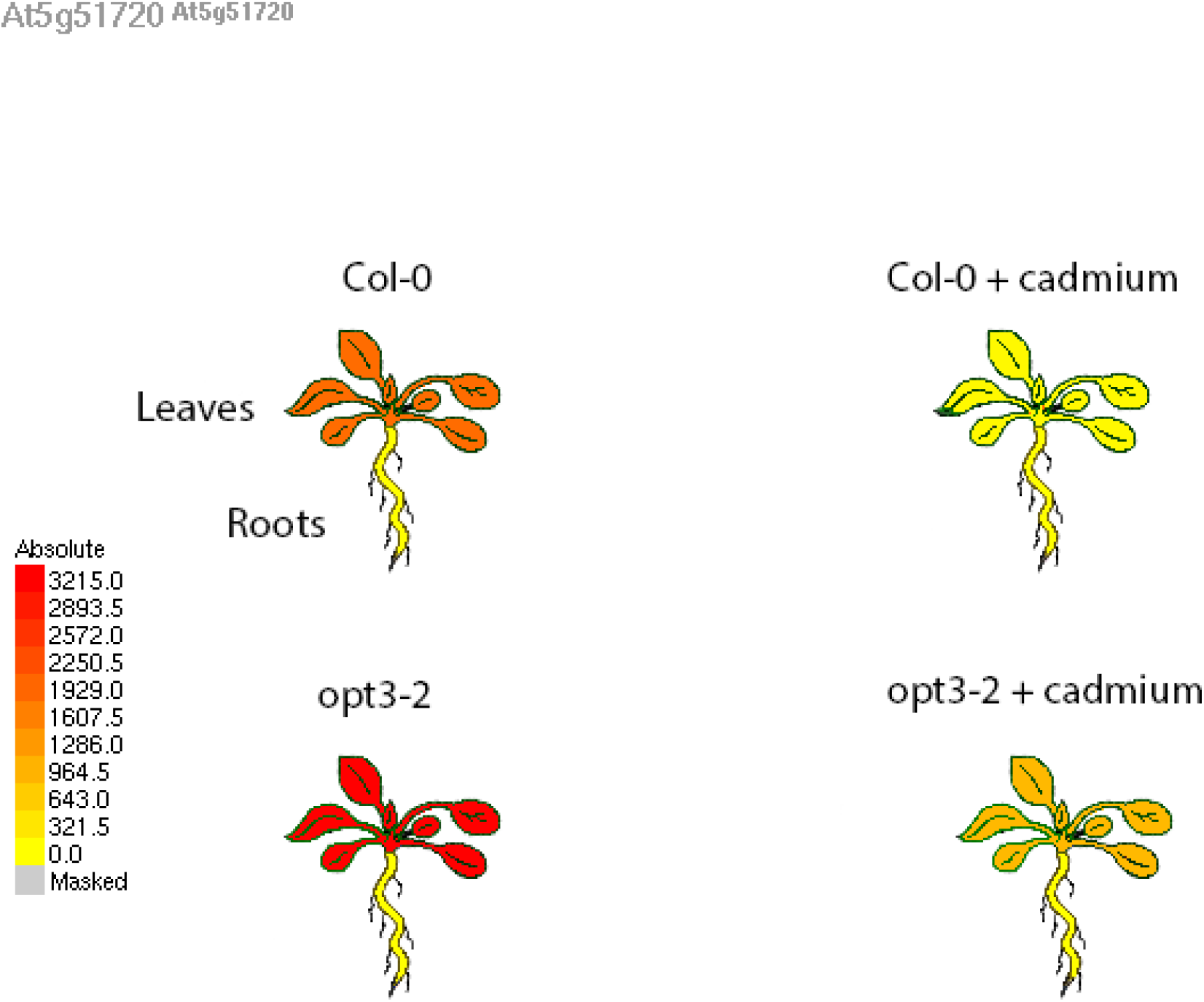
eFP browser image of RBOHD expression

**Fig. S4:**
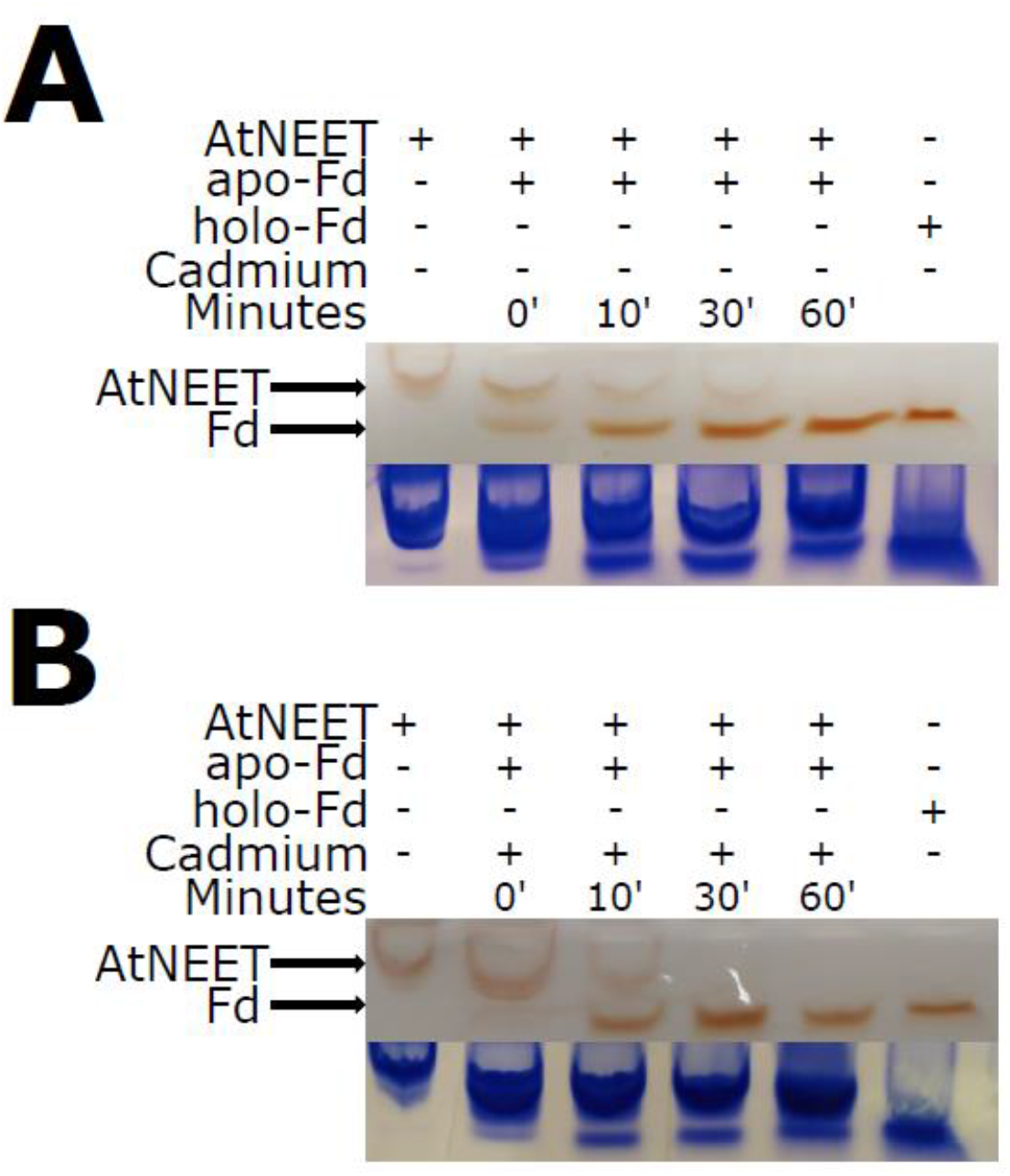
Native PAGE gel separation of AtNEET and ferredoxin (Fd). The red iron sulfur clusters transfer between AtNEET and Fd *in vitro* over time. **(A)** A normal transfer in the absence of Cd. **(B)** Impeded transfer of iron sulfur clusters in the presence of 50μM CdCl_2_

**Table S1:**

List of the genes included in the *expanded root ferrome* and *the leaf ferrome*, including gene symbols and expected sign of differential expression during Fe deficiency.

**Table S2:**

List of all genes tested, their counts per million and log_2_ fold changes between genotypes and treatments.

**Table S3:**

Fold changes and expected sign of Fe marker genes used to generate the Venn diagrams found in Fig. 2a, b.

**Table S4:**

Fold changes and counts per million of genes used to generate heatmaps found in Fig. 2c, d.

**Table S5:**

Primers used for qRT, figure S2

## Acknowledgements

This work was supported by funding from the National Science Foundation (IOS-1734145 and MCB-1818312 to DMC, NSF-BSF MCB-1936590 and IOS-1932639 to RM, and BSF 2015831 to RN), National Research Foundation, South Africa (NRF-116346 and NRF-109083 to MK,), the Bond Life Sciences Early Concept Grant (DGMC and RM), and the University of Missouri. Collaborative research between the D.M.C. and M.K. groups is funded by a DST-NRF Centre of Excellence in Food Security award (Project ID 170202).

## Author contributions

SAM, MAK, NACG conducted the RNA sequencing experiment, AG conducted the H_2_O_2_ quantification in leaves, JL conducted the H_2_O_2_ quantification in roots, RH and HHK conducted the photosynthetic efficiency experiments, HM conducted the AtNEET spectroscopy. SAM conducted the bioinformatics analysis. SAM, DMC, MAK FG, MK, HHK, RN, RM wrote the manuscript.

